# Machine Learning Ensemble Reveals Distinct Molecular Pathways of Retinal Damage in Spaceflown Mice

**DOI:** 10.64898/2026.01.22.701181

**Authors:** James A. Casaletto, Ryan T. Scott, Aahan Rathod, Aarav Jain, Aarthi Chandar, Aditi Adapala, Aditya Prajapati, Agastya Nautiyal, Anagha Jayaraman, Ananya Boddu, Anish Kelam, Anishka Jain, Bella Pham, Dhruv Shastry, Diya Narayanan, Eashan Kosaraju, Elior Paley, Fabian P. Uribe, Ibrahim Shahid, Isabel Ye, Jessica Wu, Joshua Lin, Krithikha Srinivas, MarcAnthony Paolieri Della Monica, Margaret Hitt, Matthew Lin, Maxwell Volkan, Misha Kharya, Mrinalini Kaul, Muhammad A. Jaffer, Mushtaq Ali, Naomi Z. Chang, Nishant Ashri, Noélie Boquet Couderc, Phani Paladugu, Rohin Sood, Ronak Hiremath, Rudransh Pathak, Saanvi Dogra, Samarth Srinivas, Shawnak Samaddar, Shrikar Gopinath, Shriya Sawant, Sophie Cai, Vania Pala, Vinitha Nair, Zhihan Shi, S. Anand Narayanan, Daniya Mundackal Thomas, Anna Lewkowicz, Ethan Waisberg, Joshua Ong, Samrawit Gebre, Jonathan M. Galazka, Parag A. Vaishampayan, Lauren M. Sanders, Xiao Wen Mao

## Abstract

**Background:** Spaceflight-associated neuro-ocular syndrome (SANS) poses significant risks to astronaut visual health during long-duration missions, yet its underlying molecular mechanisms remain incompletely understood. Oxidative stress and apoptosis are candidate molecular drivers, but their transcriptomic signatures in spaceflight-exposed retinal tissue have not been systematically characterized.

**Methods:** We applied a machine learning ensemble of linear regression models to predict two ocular phenotypes: 4-hydroxynonenal (4-HNE) immunostaining as a marker of lipid peroxidation-mediated oxidative damage; and TUNEL positivity as a marker of apoptotic cell death. In this observational study, we use bulk retinal gene expression data obtained from a controlled experiment with ground control and spaceflown mice to predict these phenotypes. Gene Ontology pathway enrichment was performed on the most predictive gene sets for each phenotype.

**Results:** The 4-HNE phenotype was predicted by genes that converge on membrane-associated pathways, photoreceptor protein modification, synaptic dysfunction, and extracellular matrix dysregulation, including *B2m*, *Tf*, *Cnga1*, *mt-Nd1*, *Snap25*, and *Efemp1*. The genes predicting the TUNEL phenotype revealed a distinct signature emphasizing stress-induced apoptosis, rod photoreceptor degeneration, and endoplasmic reticulum dysfunction, with *Ddit4*, *Nrl*, *Rom1*, *Reep6*, and *Gabarapl1* emerging as central regulators.

**Conclusions:** Oxidative lipid peroxidation and apoptotic cell death represent complementary and molecularly distinct pathological mechanisms in spaceflight-exposed murine retinal tissue. The gene signatures provide a putative molecular framework for developing noninvasive biomarkers and therapeutic targets to monitor and protect astronaut visual health during long-duration and deep-space missions.

## Introduction

Prolonged microgravity exposure produces multiple physiological and pathological changes within astronauts^1^, many constituting SANS. This syndrome encompasses changes in ophthalmologic tissues during long-duration spaceflight, including optic disc edema, posterior globe flattening, choroidal thickness changes, retinal folds, and oxidative damage within the blood-brain barrier^2–4^. These manifestations pose serious risks during long-term missions and represent obstacles to future deep space exploration. SANS prevalence among astronauts is high. Approximately 70 percent of ISS astronauts reported posterior eye swelling, with over 45 percent reporting near distance visual acuity changes during missions exceeding 30 days, compared to 23 percent experiencing symptoms during shorter exposures^3,4^. Structural changes, visual acuity discrepancies, and magnetic resonance imaging findings consistently demonstrate SANS pathology^2,4,5^. There are several hypothesized pathways involved with crew-members developing SANS including cephalad fluid shifts and elevation of intracranial pressure due to microgravity, retinal ischemia, oxidative stress from microgravity and radiation exposure, ocular and brain structural and functional changes including optic nerve sheath distension, specific responses to spaceflight with certain biochemical pathways (e.g. folate and vitamin-B12 dependent one-carbon metabolism pathways), as well as sex, age, high-salt diet, and cardiovascular health influence developing of SANS with astronauts^6^.

Microgravity-induced cephalad fluid shifts represent a key mechanism underlying SANS pathophysiology^4,5^. During ascent, cephalic, cervical, and pectoral tissues swell while femoral and abdominal regions slim. Upon reaching orbit, fluids achieve homeostatic distribution maintained throughout missions^5^. This redistribution triggers physiological adaptations in response to the spaceflight environment, including cardiovascular adaptations, changes in blood circulation, vasculature, and headward fluid shifts in response to space travel, leading to decreases in venous pressures, loss of plasma volume, and orthostatic intolerance. More specifically, venous congestion increases ocular venous pressure, contributing to choroidal expansion through vortex vein drainage lacking autoregulatory control^5,7^. Choroidal expansion also occurs during head-down tilt parabolic flights^4,5^. Cephalad fluid shift causes optic disc edema and contributes to optic disc protrusion and osteoma-like formations^2,4^. These fluid shifts also lead to ocular structural changes, namely ocular and cerebral blood brain barrier adaptations^8,9^.

Beyond hemodynamics, microgravity and space radiation elicit retinal oxidative stress and apoptosis underlying SANS alterations. Murine studies demonstrate elevated 4-HNE (due to lipid peroxidation) and increased TUNEL-positive nuclei (due to apoptotic cell death) in the inner nuclear layer (INL) and ganglion cell layer (GCL) following spaceflight and simulated radiation^3^, with additional evidence of adverse photoreceptor and retinal functional changes^10^. Various studies have shown that crew members develop increased oxidative stress in response to the spaceflight environment, influencing bone loss, cardiovascular function, immune insufficiency, metabolic dysfunction, neurological impairment, among other adaptations^11^. Reactive oxygen species (ROS) trigger lipid peroxidation producing electrophilic aldehydes like 4-HNE. Caspase activation drives DNA fragmentation detectable through the TUNEL assay^12^.

There have been various studies exploring spaceflight associated neuro-ocular adaptations. These include changes in hypovolemia, cardiac atrophy, region-specific vascular remodeling, sensory-neural adaptation; in particular, are the ocular and neural structural adaptations^13^. It has been reported that after 6 months of spaceflight on the International Space Station, astronauts had ophthalmic abnormalities consisting of optic disk edema, posterior globe flattening, choroidal folds, cotton wool spots, retinal nerve fiber layer thickening, and decreased near vision and hyperopic shifts, with suggestions of elevated intracranial pressure^2^. Moreover, a postflight survey of astronauts has shown that 29 and 60% of astronauts on short-duration and long-duration missions, respectively, experienced degradation in distant- and near-vision visual acuity. Brain structural adaptations are also observable, with crowding of the brain sulci and brain ventricular volume changes^14,15^.

These observations can be more comprehensively studied through rodent research. Indeed, Overbey et al investigated retinal RNA transcriptomic data of male mice on a 35-day International Space Station (ISS) mission which revealed over 600 differentially expressed retinal genes which enrich for visual perception, phototransduction, and photoreceptor phenotypes, retinitis pigmentosa, diabetic retinopathy, macular degeneration, and retinal detachment^16^. 4-HNE staining of the eye was conducted, which showed increased cone photoreceptors, INL, and GCL post-spaceflight ^17^. TUNEL assays were also conducted which show evidence of space-induced oxidative damage from mitochondrial apoptosis^3^. However, molecular pathways linking transcriptomic remodeling to oxidative injury and apoptosis within SANS remains undefined. Previous studies such as this treated RNA sequencing, histology, and imaging as parallel endpoints without identifying quantitative linkage between gene expression patterns and phenotypic assays^3^. To address this gap, our observational study applied a multi-modal machine learning ensemble that integrates the retinal transcriptomic profiles with quantitative bioimaging phenotypes (4-HNE immunoreactivity and TUNEL labeling) from spaceflight-exposed and ground control samples in the study from Overbey et al to identify consensus gene signatures predictive of tissue-level oxidative injury and apoptosis.

## Methods

We developed a multi-modal machine learning pipeline to identify genes whose expression profiles predict quantitative bioimaging phenotypes of retinal oxidative stress and apoptosis. Figure 1 summarizes the complete workflow; the following subsections detail each stage.

**Figure 1.**
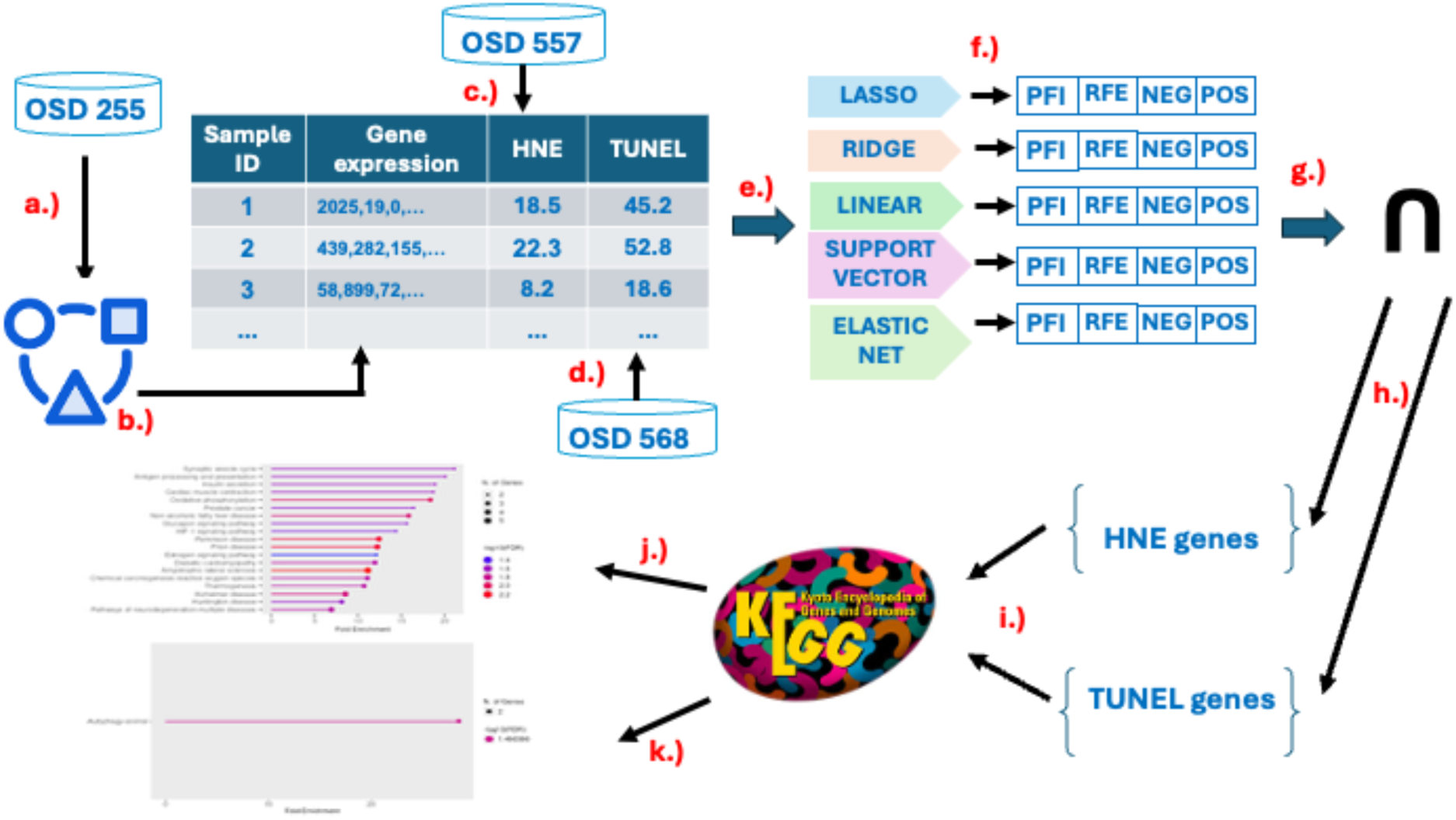
Machine learning pipeline. a.) Leveraged RNA-sequencing gene expression data from OSD-255. b.) Applied filters and data transformations to gene expression data to prepare for training models. c.) Assigned 4-HNE phenotype values and gene expression data for each sample. d.) Assigned TUNEL phenotype values and gene expression data for each sample. e.) Trained ensemble of linear regression models on filtered and transformed gene expression data to predict phenotype values. f.) Identified genes most predictive of phenotype values for each model using permutation feature importance (PFI), recursive feature elimination (RFE), largest positive coefficients, and largest negative coefficients. g.) Found the intersecting majority consensus of most predictive genes across all models in ensemble. h.) Identified genes uniquely predictive of TUNEL and 4-HNE phenotypes and i.) sent them to the KEGG database to obtain enriched gene sets for the j.) 4-HNE and k.) TUNEL experiments.

### Data

The Rodent Research 9 (RR-9) mission, conducted by NASA in August 2017, investigated these effects using murine model organisms, with initial characterization demonstrating blood-retinal barrier disruption and ocular adaptations^8^. Two datasets from this mission were collected and curated by the NASA Ames Life Sciences Data Archive^18^, and one dataset was curated by NASA GeneLab, all three now accessible through NASA’s Open Science Data Repository^19^.

OSD-557 and OSD-568 contain bioimaging data which explore the retinal consequences of spaceflight using 4-HNE immunofluorescence to assess oxidative stress and TUNEL bioimage staining to detect apoptosis, respectively^16,20^. These studies aimed to elucidate molecular and cellular mechanisms underlying SANS to inform potential therapeutic strategies. Complementing the imaging data, OSD-255 provides bulk RNA sequencing gene expression profiles from retinal tissue^16^. There were 16 male C57BL/6J 10-week-old mice in this experiment. Eight of the mice were flown in space for 35 days with a 12-hour light/dark cycle. They were fed nutrient-upgraded rodent food bars *ad libitum*. Within 38 hours of splashdown, all flight mice were euthanized, and their eyes were removed and prepared for analysis. The retinas from the left eyes were fixed in 4% paraformaldehyde in phosphate-buffered saline (PBS) for immunohistochemical staining. The right eyes were snap-frozen using liquid nitrogen for downstream transcriptomic analysis. The eight habitat ground control mice were otherwise exposed to identical conditions.

In OSD-255, RNA-seq was performed using single-stranded, ribo-depleted sequencing with an External RNA Controls Consortium (ERCC) spike-in mix on an Illumina HiSeq 4000. The sequencing data were trimmed using TrimGalore and aligned using both RNA-seq by Expectation-Maximization (RSEM) and Splice Transcripts Alignment to a Reference (STAR) software^21,22^. RSEM and STAR are included in the GeneLab transcriptomic processing pipeline^23^. We merged both gene expression data files (unnormalized STAR counts and unnormalized RSEM counts) as predictors in our model. In OSD-557, oxidative stress was evaluated using 4-HNE immunostaining, a marker of lipid peroxidation. ImageJ analysis quantified 4-HNE-positive cells and immunofluorescence intensity in the cone photoreceptors, INL, and GCL. Key metrics assessed included total 4-HNE-positive cells (`sumcount`), 4-HNE-positive endothelial cells (`sumEC`), and endothelial cell density (`denEC`). For OSD-568, TUNEL assays were used to label DNA fragmentation, a hallmark of apoptosis, in stained retinal sections from spaceflight and ground control mice. The CellProfiler software package quantified TUNEL-positive nuclei, focusing on the GCL, INL, and outer nuclear layer (ONL), to provide insight into regional apoptotic activity. For our machine learning models, we isolated the endothelial cell-specific phenotype variables (total EC is `sumEC` and EC density is `Density_EC`) since they directly align with the study’s focus on how stress on retinal capillaries can compromise the blood-retinal barrier (BRB) and lead to retinal degeneration.

Figure 2 shows the box-and-whisker plots of the distributions of 4-HNE `sumEC` and TUNEL `Density_EC` phenotype measurements in the ground control and spaceflight groups. Using the Student’s t-test, we found that the difference between the means of the ground control and spaceflight distributions for both phenotypes is statistically significant (p<0.05).

**Figure 2.**
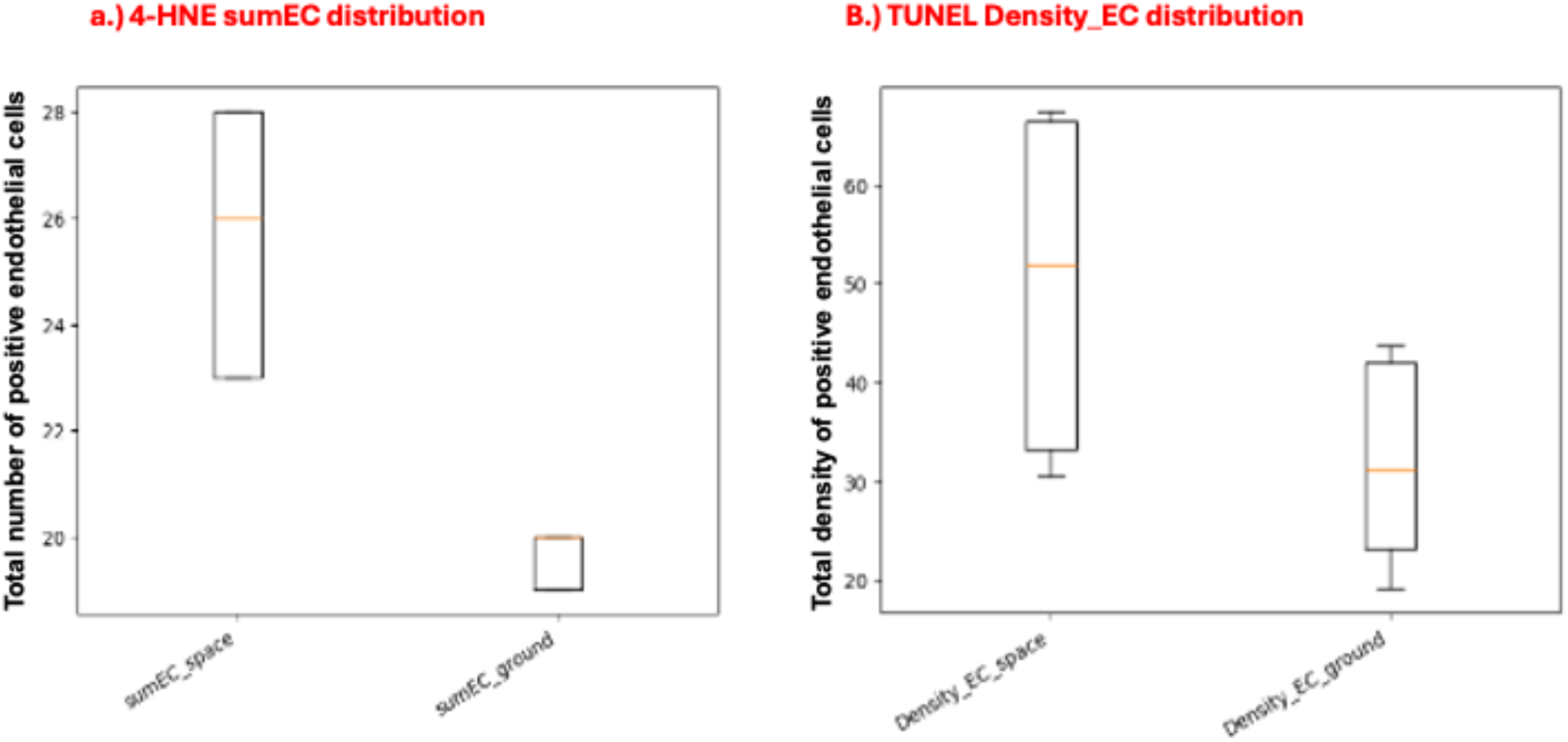
a.) Distributions of 4-HNE phenotype measurements (`sumEC`) on a scale from 0 to 30 for spaceflown mice and ground control mice. b.) Distributions of TUNEL phenotype measurements (`Density_EC`) on a scale from 0 to 70 for spaceflown and ground control mice.

### Data Preparation

We merged both the RSEM and STAR abundance files as a form of data augmentation to help offset the high dimensionality of our gene expression data. We then applied various gene filtering methods to reduce the dimensionality of the feature space. First, we removed genes that did not have counts for all samples. Next, we removed low-abundance genes from the data with a threshold of 50 or fewer transcripts in 80% or more of the samples. Third, we considered only protein-coding genes. Finally, we kept the 1,000 genes most highly correlated to the phenotypic target. Table 1 depicts the number of genes removed by each filter.

**Table 1.** Filters applied to the data to reduce dimensionality, beginning with 56,840 genes and ending with 1,000 genes for all the experiments.

| Filter name | Number of genes before applying filter | Number of genes removed by filter | Number of genes remaining after filter |
| --- | --- | --- | --- |
| Filter not-available | 56,840 | 33,421 | 23,419 |
| Filter low-abundance | 23,419 | 12,477 | 10,942 |
| Filter non-coding | 10,942 | 553 | 10,389 |
| Filter non-correlated | 10,389 | 9,389 | 1,000 |

In addition to filtering the data, we performed the following data transformations on the gene expression data before training the model. First, we normalized the data using transcripts per million (TPM) to account for variable sequencing depth and gene lengths. Second, we scaled the data to z-scores so that we could use coefficient-based feature importance methods.

The PCA plots of Figure 3 show clearly separated sample gene expression clusters for high and low values of the 4-HNE and TUNEL phenotypes. This separability in reduced feature space suggests that gene expression may be used to predict the phenotype values for each assay.

**Figure 3.**
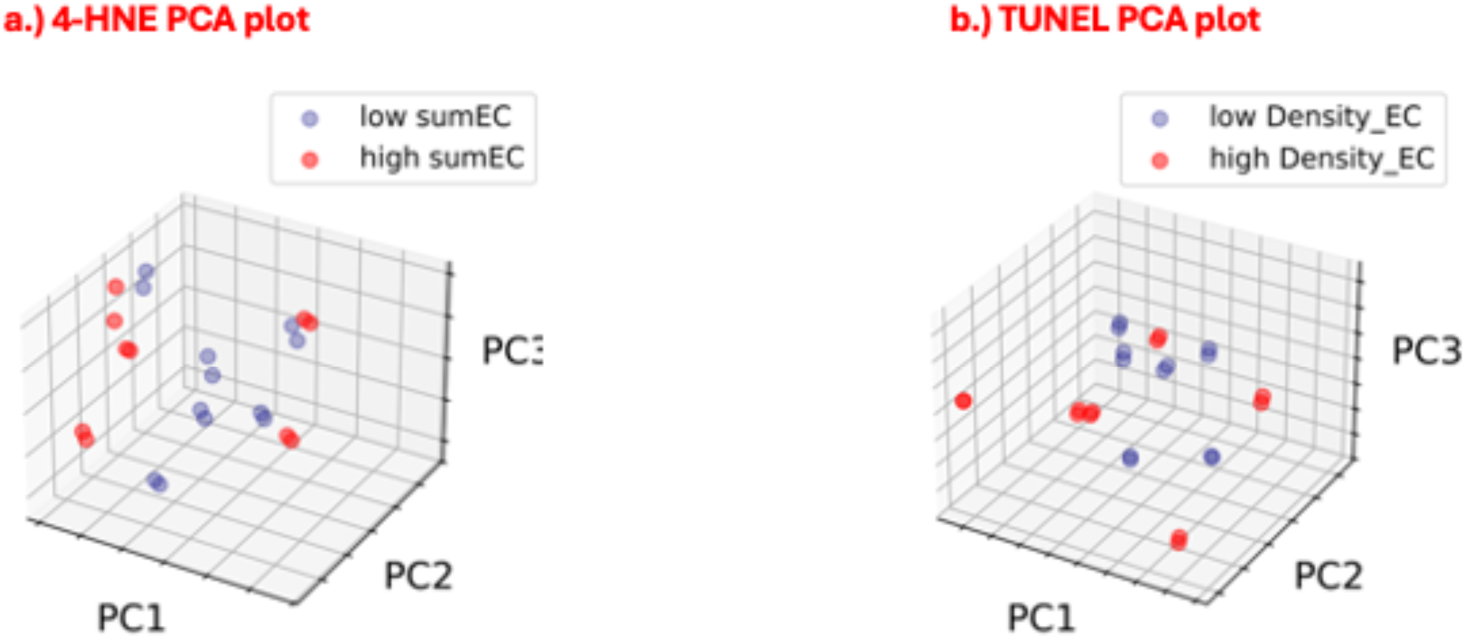
PCA plots of gene expression data colorized by low and high phenotype values for a.) 4-HNE `sumEC` and b.) TUNEL `Density_EC`. Each sample has 2 closely paired gene expression values (one from STAR protocol and one from the RSEM protocol).

### Machine Learning Ensemble

To investigate whether spaceflight-induced retinal stress can be detected directly from transcriptomic signatures, we built a multi-modal machine-learning ensemble to predict 4-HNE and TUNEL phenotypes from gene expression profiles. The goal of these experiments is to determine whether transcriptomic patterns could reliably predict oxidative damage (4-HNE) and apoptotic activity (TUNEL). The use of multiple linear regression models captures linear signatures of retinal injury induced by spaceflight. By combining models that each learn biological signals in different ways, the ensemble provides a more comprehensive and robust estimate of gene-phenotype relationships than any single method alone. We adapted the ensemble machine learning methodology of Casaletto et al.^24^, which employed multiple linear classifiers and majority voting to identify consensus gene sets predictive of spaceflight-induced phenotypic changes.

We chose 5 different machine learning models to form our ensemble: linear regression, elastic net, support vector regression, ridge regression, and lasso regression. The basic form of linear regression estimates a continuous response from a linear combination of predictor variables, through estimating coefficients by minimizing the sum of squared residuals between observed and predicted values^25^. Support vector regression uses a kernel-based approach that fits a function within a margin of error and maximizes the margin around it to show linearly or nonlinearly separable relationships between the inputs and the target, depending on the choice of kernel^26^. Ridge regression adds an L2 penalty (proportional to the squared magnitude of the coefficients) to the least squares loss, which shrinks coefficient magnitudes towards zero. This reduces multicollinearity and overfitting in high-dimensional settings^27^. Lasso regression adds an L1 penalty on coefficient magnitudes, driving many coefficients to exactly zero, which effectively performs feature selection and identifies the most predictive variables in the dataset^28^. Elastic net regression combines the penalties of both L1 (lasso) and L2 (ridge) regularization so that it shrinks coefficients and performs variable selection. This is especially useful when predictors are highly correlated^29^.

Using an ensemble of regression models instead of relying on a single model provides the benefit leveraging the unique strengths of each algorithm and reduce model-specific bias and variance, particularly with a high-dimensional feature space such as gene expression data. Each regression method learns patterns in its own way. For example, regularized models like ridge, lasso, and elastic net limit the influence of weak or noisy genes, whereas support vector regression uses kernel functions to define boundary relationships. Together, the ensemble can model both simple and complex gene-phenotype patterns. Ensemble learning often improves reliability in small, high-dimensional biological datasets where no single model performs best across all conditions. All five models were built using `scikit-learn` version 1.7.2.

### Model Selection

We used a train/test split of 70/30 and validated using k-fold cross validation across the training data. This ensures that each sample is used for both training and validating and reduces overfitting. It also has the effect of giving a more stable estimate of performance^30^. Because our dataset is so small (n=16), we used the leave-p-out (p=2) method of cross-validation for model selection. This method is ideal for small datasets because it makes efficient use of limited samples and provides a more stable estimate of model performance. To assess the quality and generalization capacity of the regression models, we calculated the R^2^ (coefficient of determination). This metric, calculated with the `scikit-learn` library, provides an evaluation of the models’ explanatory power and absolute magnitude of prediction error. The R^2^ score is a measurement that quantifies the proportion of the variance in the dependent variable (the phenotype values) that the model predicts given the independent variables (the gene expression profiles). The R^2^ score serves as a relative measure of model fit, comparing the model’s performance to a baseline prediction of the mean of the observed data. A score of 1.0 indicates a perfect prediction, with the model explaining 100% of the variability in the target data. A score of 0.0 indicates the model performs no better than the baseline prediction of mean target value, and a negative score indicates the model performs worse than simply predicting the mean of the target values. We selected the models with an R^2^ score of 0.9 or higher as a conservative threshold based on an informal assessment of R^2^ score thresholds defined in the scientific literature. After training each model, we identified the features most predictive of the phenotype values on the held-out test data using the feature-selection methods described below.

### Feature Selection

We used two approaches from the `scikit-learn` package for feature selection using the test data set: PFI and RFE. The PFI approach randomly permutes the values of each feature across the test samples one by one and calculates the model performance before and after having done so, which measures how much each feature affects model performance^31^. The degree to which the model performance changes is assigned to each feature as its importance metric, with larger degradations indicating greater importance. RFE, by contrast, repeatedly removes the least predictive features from the model through an iterative retraining process, revealing which features are most important to the model’s internal predictive structure. The algorithm continues rebuilding models and removing features until the desired number of features remain^32^. Using both PFI and RFE allows us to capture complementary information: PFI reflects a feature’s external impact on model performance; and RFE identifies features that the model internally depends on most. For both feature selection techniques, we used the coefficient of determination scoring metric R^2^. We also used coefficient magnitude to identify the features with the largest coefficients as a proxy for feature importance. We selected the 40 largest magnitude positive coefficients and the 40 largest magnitude negative coefficients. In summary, each ML algorithm generates 4 lists of 40 genes: genes ranked by PFI; genes ranked by RFE; genes associated with the 40 largest positive coefficients; and the genes associated with the 40 largest negative coefficients.

We intersect the PFI and RFE gene sets, then intersect that result with the union of the positive and negative coefficient sets. Positive coefficients were interpreted as features whose higher values are associated with an increase in the predicted outcome (assumed to be “up-influencing” the response), while negative coefficients indicate features whose higher values are associated with a decrease in the response (assumed to be “down-influencing” the response). Intersecting these lists increases stability and reproducibility by ensuring a gene must be consistently important across multiple, independent criteria rather than appearing (due to noise) in any single method. We keep the genes that appear in the list of the simple majority of the models whose R^2^ performance is higher than 0.9 to define the final list of genes for the ensemble, only keeping genes that are repeatedly selected by high-performing models in the final ensemble list. This cross-model comparison reduces model-specific bias and noise and emphasizes features that are stable and robustly predictive across methods.

### Gene Set Enrichment Analysis

Our bioinformatic pipeline concludes with gene set over-representation, enrichment, and pathway analysis. Predictive genes from machine learning models underwent gene set enrichment analysis using the online Web portal for the Molecular Signatures Database (MSigDB) against the Gene Ontology collections (biological process, molecular function, cellular component) and Human Phenotype Ontology sets. MSigDB enrichment tests whether predictive genes overlap significantly within curated gene sets representing pathways, functions, processes, and phenotypes^33^. This overrepresentation test counts overlap between our lists and curated sets, computes p-values, and applies multiple test correction (FDR < 0.05). No enrichment score thresholds were applied.

### Comparison to causal inference ensemble

As a form of validation for our approach, we compare the results of our machine learning ensemble to the results from a known *in silico* causal inference platform called the Causal Research and Inference Search Platform (CRISP). CRISP itself is an ensemble of 4 machine learning algorithms which leverages the concept of invariance as an *in silico* proxy for causality. Casaletto et al used the CRISP platform to identify genes putatively causal of non-alcoholic fatty liver disease in spaceflown murine liver tissue^34^.

## Results

In this section, we discuss the training and testing performance of each of the models in the ensemble. We then describe the genes that our ensemble identified as most predictive of the 4-HNE and TUNEL phenotypes. Finally, we discuss the gene sets that are most enriched by the most predictive genes for the 4-HNE phenotype, TUNEL phenotype, and for both phenotypes.

### 4-HNE model performance

Figure 4 shows the distributions of cross-validation training and held-out test R^2^ scores for the 4-HNE phenotype. We include the most predictive genes only from those models whose held-out test R^2^ scores are 0.9 or higher and then take the genes that appear in a simple majority of those high-performing models as our final gene sets.

**Figure 4.**
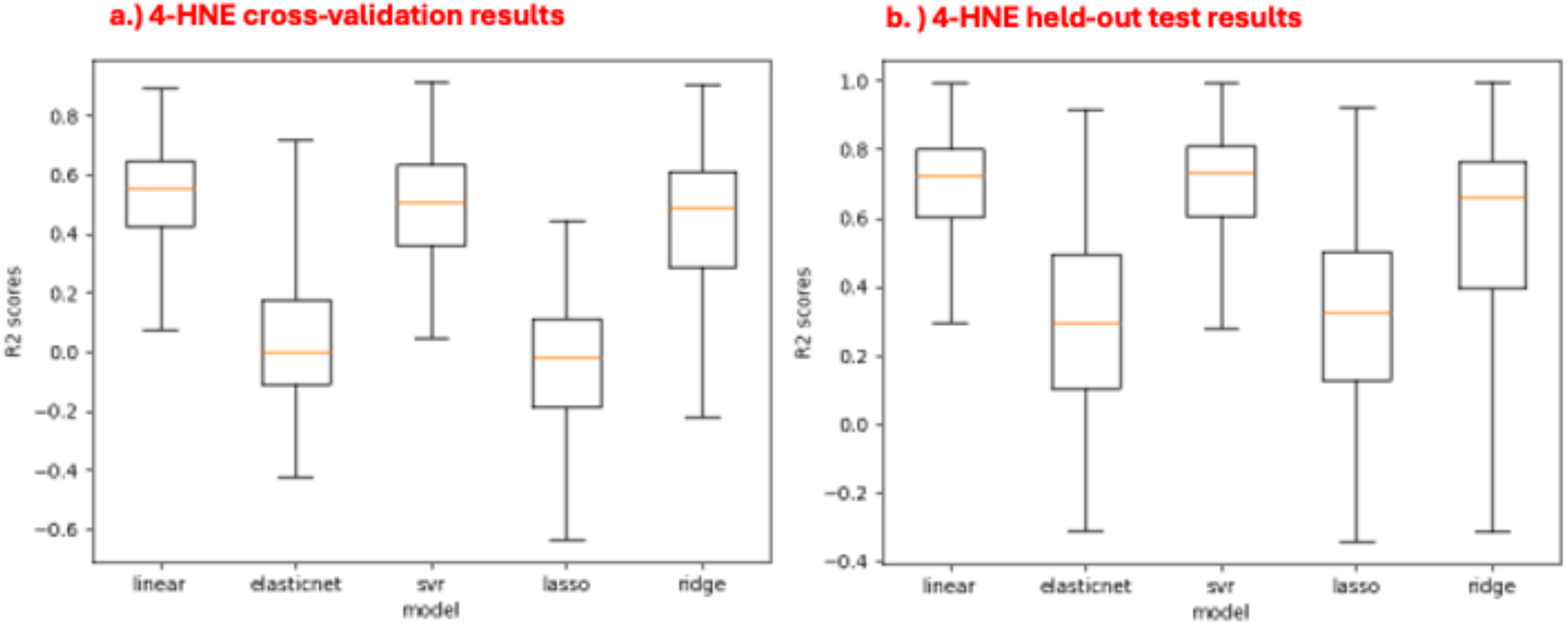
Performance results from the a.) cross-validation set and b.) held-out test set for each regression model predicting the 4-HNE phenotype value.

In the case of 4-HNE, three of the five models achieved R^2^ scores higher than 0.9 (linear regression, lasso regression, and ridge regression). In the construction of the final gene set for 4-HNE, we select those genes which appear in the majority (2 or 3) of feature importance lists of these 3 algorithms. As shown in Figure A1 of the Appendix, the genes most predictive of the 4-HNE phenotype values are approximately uniformly distributed across the background of all genes, demonstrating that the results are not biased due to the homoscedastic nature of transcriptomic count data.

### TUNEL model performance

Figure 5 shows the distributions of cross-validation training and held-out test R^2^ scores for the TUNEL phenotype. Again, we include the most predictive genes only from those models whose held-out test R^2^ scores are 0.9 or higher and then take the genes that appear in a simple majority of those high-performing models as our final gene sets.

**Figure 5.**
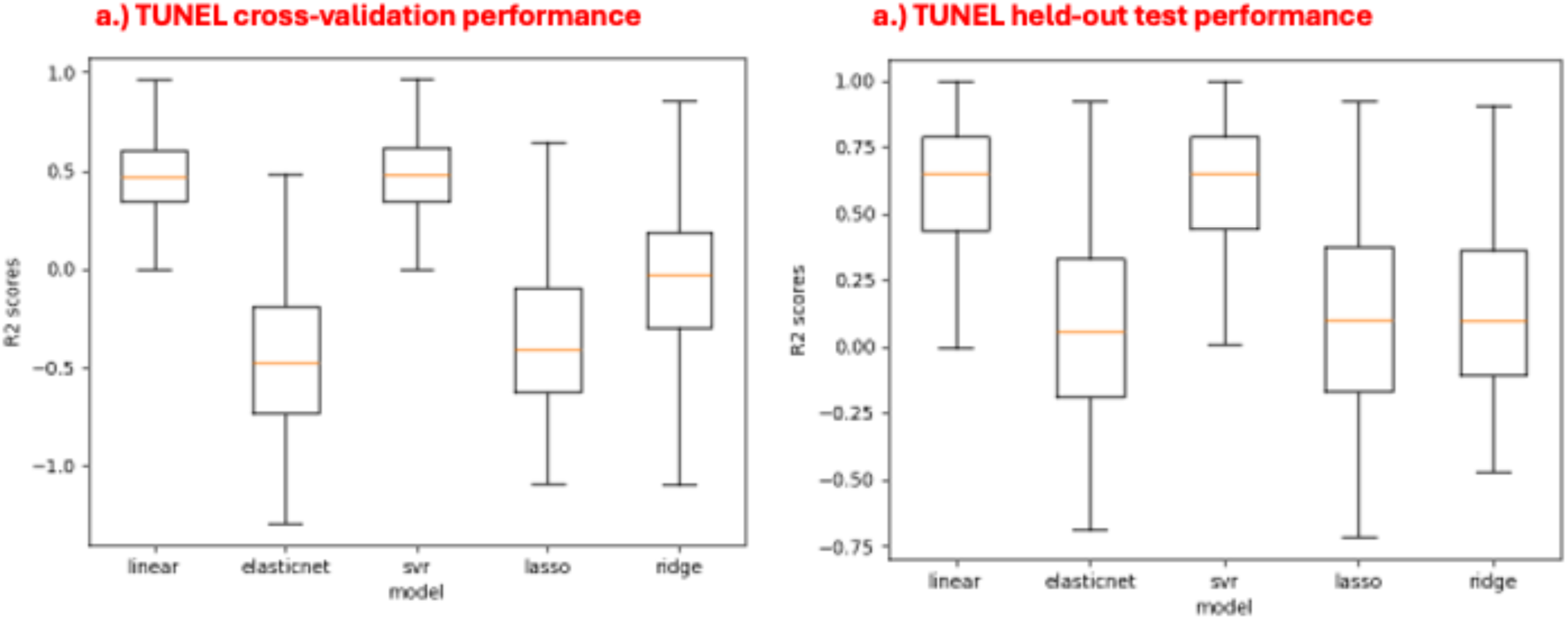
Performance results from the a.) cross-validation set and b.) held-out test set for each regression model predicting the TUNEL phenotype value.

Similarly, in the case of TUNEL, three of the five models achieved R^2^ scores higher than 0.9 (linear regression, elastic net regression, and support vector regression). In the construction of the final gene set for TUNEL, we select those genes which appear in the majority (2 or 3) of feature importance lists of these 3 algorithms. As shown in Figure A2 of the Appendix, the genes most predictive of the TUNEL phenotype values are approximately uniformly distributed across the background of all genes, demonstrating that the results are not biased due to the homoscedastic nature of transcriptomic count data.

The model performance metrics revealed strong predictive ability on both 4-HNE and TUNEL phenotypes. The linear regression and support vector regression models consistently outperformed the other models on both cross-validation training and held-out test sets and have similar results to each other. The elastic net and lasso models consistently performed the worst across the datasets and have similar results to each other. The ridge regression model performance was approximately average across all 5 algorithms and both cross-validation training and held-out test datasets. These results suggest our models are not overfit (the cross-validation training and held-out test scores are similar), that our ensemble provides model variance, and that our sample size is likely adequate relative to the number of features we’re using, though caution is still warranted due to our small sample size of n=16. The fact that our unregularized linear regression model outperforms the regularized models (lasso, ridge, elastic net) in the held-out test data suggests that we may be over-penalizing those regularized models for their complexity (though we used grid search to identify the optimal value for the alpha regularization parameter). This in turn suggests that most genes in the feature space are informative, validating our choices in the filtering preprocessing step.

### Genes uniquely predictive of the 4-HNE oxidative stress phenotype

All genes in the majority of high-performing models identified as most predictive of the 4-HNE oxidative stress phenotype are shown in Table 2. Positive average coefficient values indicate a positive correlation with the 4-HNE target, and negative average coefficient values indicate a negative correlation with the 4-HNE target.

**Table 2.**
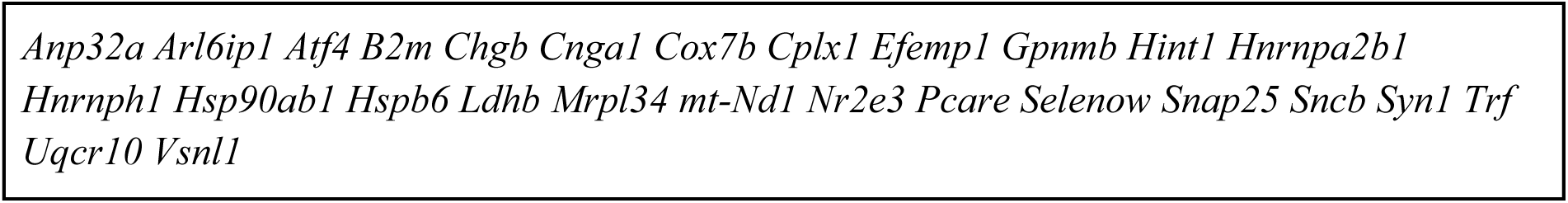
Union of sets of most predictive genes from the majority of high-performing models in predicting 4-HNE listed in alphabetical order.

Table 3 shows the functional enrichment (of the genes from Table 2) in gene sets from KEGG Gene Ontology and Human Phenotype Ontology databases. The final gene sets associated with predicting the 4-HNE phenotype involve a complex stress response encompassing structural and morphological abnormalities (e.g. optic disc pallor), photoreceptor and retinal degeneration (e.g. retinal dystrophy), and fundamental cellular processes (e.g. neuron projection). Together, this demonstrates that 4-HNE acts as a molecular bridge connecting oxidative stress to the structural and functional pathologies observed in SANS.

**Table 3.**
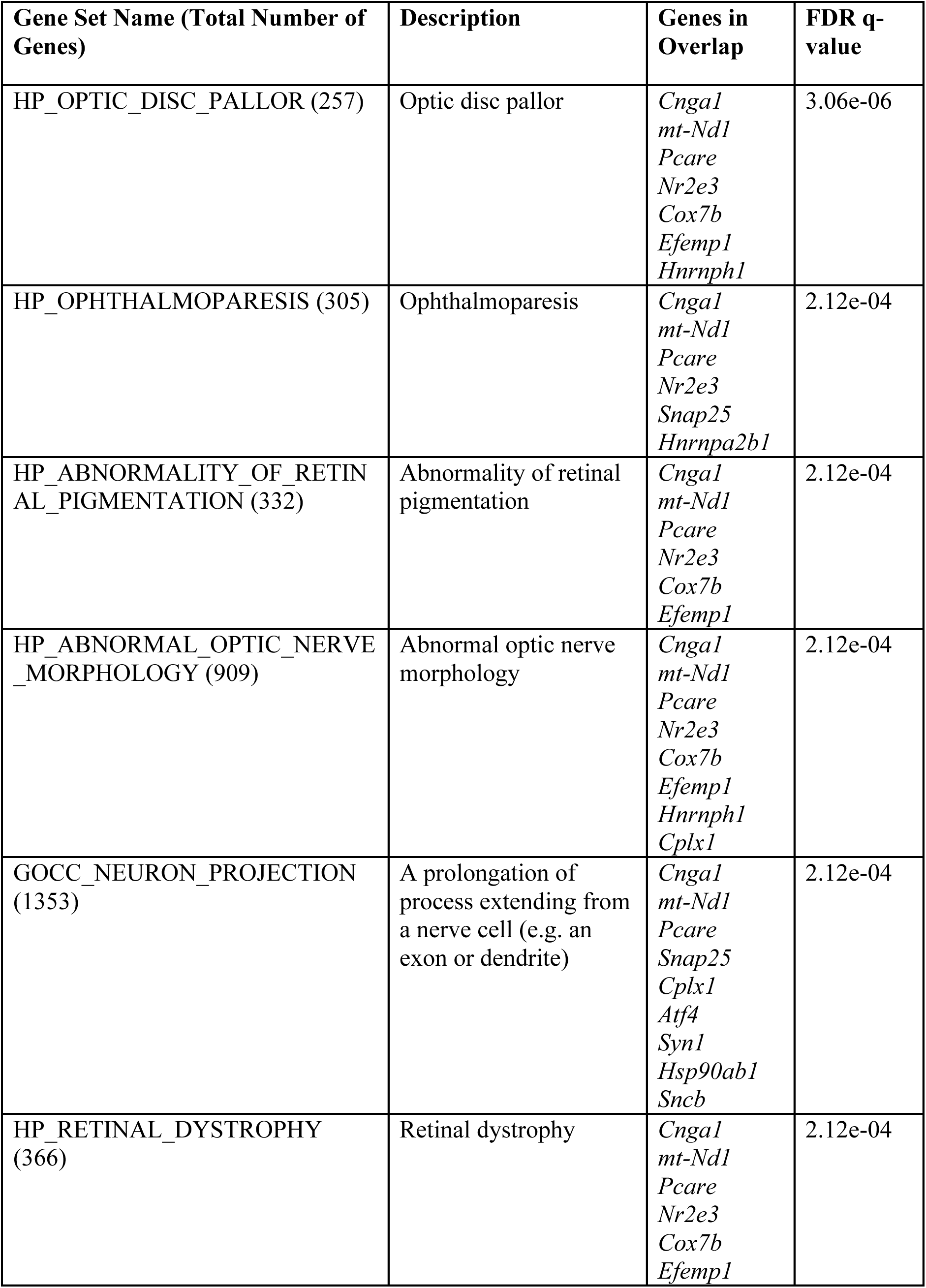
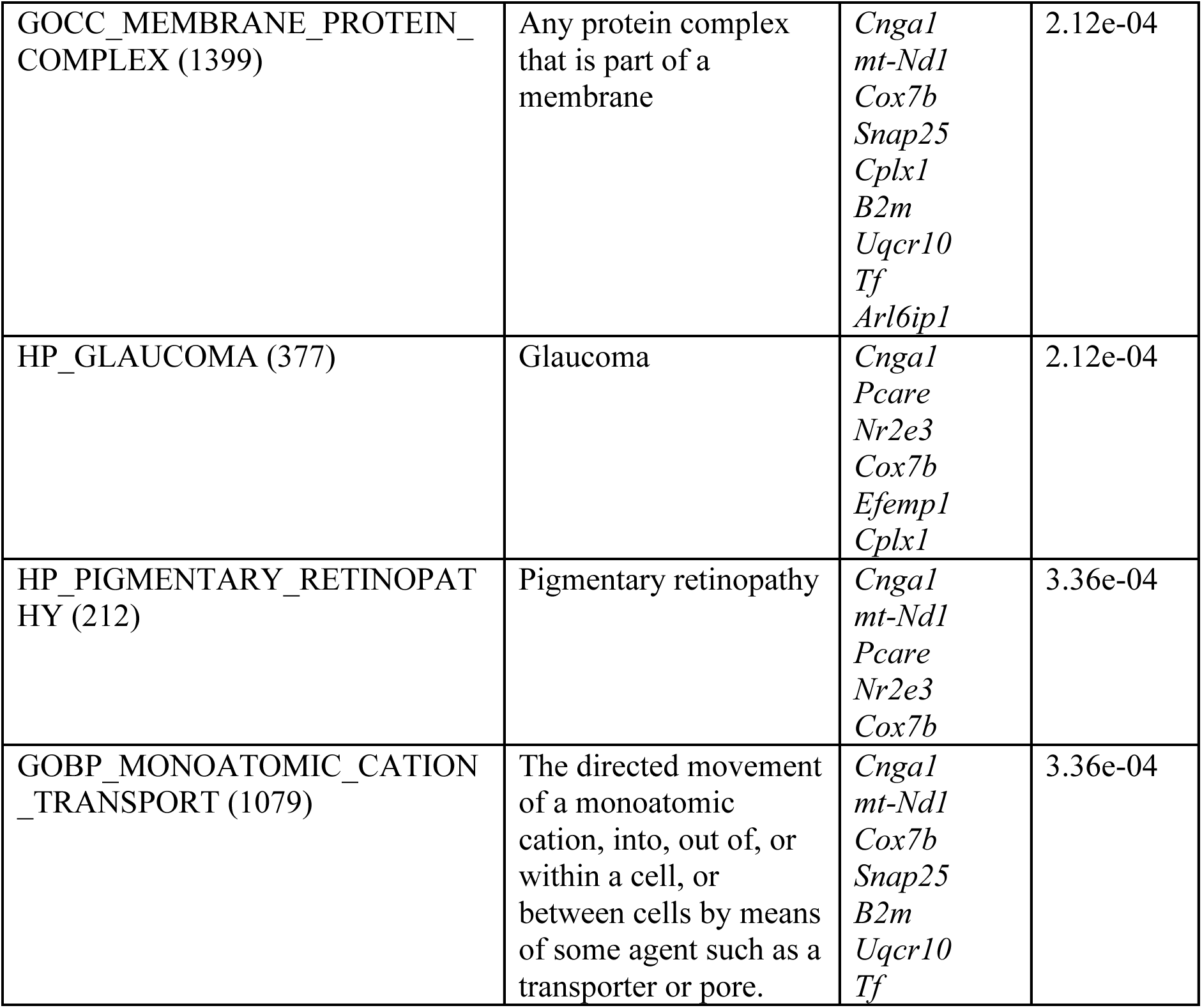
shows the top 10 pathways and gene sets significantly enriched by the genes most predictive of 4-HNE oxidative stress phenotype, using KEGG and Gene Ontology databases.

| Gene Set Name (Total Number of Genes) | Description | Genes in Overlap | FDR q-value |
| --- | --- | --- | --- |
| HP_OPTIC_DISC_PALLOR (257) | Optic disc pallor | *Cnga1*<br>*mt-Nd1*<br>*Pcare*<br>*Nr2e3*<br>*Cox7b*<br>*Efemp1*<br>*Hnrnph1* | 3.06e-06 |
| HP_OPTHALMOPARESIS (305) | Ophthalmoparesis | *Cnga1*<br>*mt-Nd1*<br>*Pcare*<br>*Nr2e3*<br>*Snap25*<br>*Hnrnpa2b1* | 2.12e-04 |
| HP_ABNORMALITY_OF_RETINAL_PIGMENTATION (332) | Abnormality of retinal pigmentation | *Cnga1*<br>*mt-Nd1*<br>*Pcare*<br>*Nr2e3*<br>*Cox7b*<br>*Efemp1* | 2.12e-04 |
| HP_ABNORMAL_OPTIC_NERVE_MORPHOLOGY (909) | Abnormal optic nerve morphology | *Cnga1*<br>*mt-Nd1*<br>*Pcare*<br>*Nr2e3*<br>*Cox7b*<br>*Efemp1*<br>*Hnrnph1*<br>*Cplx1* | 2.12e-04 |
| GOCC_NEURON_PROJECTION (1353) | A prolongation of process extending from a nerve cell (e.g. an axon or dendrite) | *Cnga1*<br>*mt-Nd1*<br>*Pcare*<br>*Snap25*<br>*Cplx1*<br>*Atf4*<br>*Syn1*<br>*Hsp90ab1*<br>*Sncb* | 2.12e-04 |
| HP_RETINAL_DYSTROPHY (366) | Retinal dystrophy | *Cnga1*<br>*mt-Nd1*<br>*Pcare*<br>*Nr2e3*<br>*Cox7b*<br>*Efemp1* | 2.12e-04 |
| GOCC_MEMBRANE_PROTEIN_COMPLEX (1399) | Any protein complex that is part of a membrane | *Cnga1*<br>*mt-Nd1*<br>*Cox7b*<br>*Snap25*<br>*Cplx1*<br>*B2m*<br>*Uqcr10*<br>*Tf*<br>*Arl6ip1* | 2.12e-04 |
| HP_GLAUCOMA (377) | Glaucoma | *Cnga1*<br>*Pcare*<br>*Nr2e3*<br>*Cox7b*<br>*Efemp1*<br>*Cplx1* | 2.12e-04 |
| HP_PIGMENTARY_RETINOPATHY (212) | Pigmentary retinopathy | *Cnga1*<br>*mt-Nd1*<br>*Pcare*<br>*Nr2e3*<br>*Cox7b* | 3.36e-04 |
| GOBP_MONOATOMIC_CATION_TRANSPORT (1079) | The directed movement of a monoatomic cation, into, out of, or within a cell, or between cells by means of some agent such as a transporter or pore. | *Cnga1*<br>*mt-Nd1*<br>*Cox7b*<br>*Snap25*<br>*B2m*<br>*Uqcr10*<br>*Tf* | 3.36e-04 |

### Comparison to results from CRISP ensemble for 4-HNE

Because CRISP is a binary classifier and 4-HNE values are continuous, we binned the 4-HNE phenotype values as “low” and “high” based on their being lower or higher than the median 4-HNE value, respectively. The top 20 genes most predictive of the 4-HNE phenotype according to the CRISP ensemble that were also identified by our approach include *Hsp90ab1* and *mt-Nd1*.

Moreover, when submitting the 20 CRISP genes to MSigDB, the gene sets that intersect with those enriched by our approach include GOCC_NEURON_PROJECTION and HP_OPTIC_DISC_PALLOR.

### Genes uniquely predictive of the TUNEL apoptosis phenotype

All genes in the majority of high-performing models identified as most predictive of the TUNEL apoptosis phenotype are shown in Table 4.

**Table 4.** Union of sets of most predictive genes from the majority of high-performing models in predicting TUNEL listed in alphabetical order.

|  |
| --- |
| *Ddit4* *Gabarapl1* *Ip6k2* *Mgarp* *Nrl* *Pdzph1* *Reep6* *Rom1* *Slc24a1* *Tcf20* *Vtn* |

The top genes most predictive of the TUNEL apoptosis phenotype (from Table 4) significantly enrich the gene sets listed in Table 5. These pathways contain genes involved in photoreceptor-specific cell death (e.g. nyctalopia), structural deterioration from cell loss (e.g. retinal atrophy), and electrophysiological dysfunction (e.g. abnormal visual electrophysiology). Unlike 4-HNE’s functional disruption of existing cellular machinery, TUNEL-positive signatures indicate the irreversible cellular commitment to cell death that characterize advanced stages of SANS pathology.

**Table 5.**
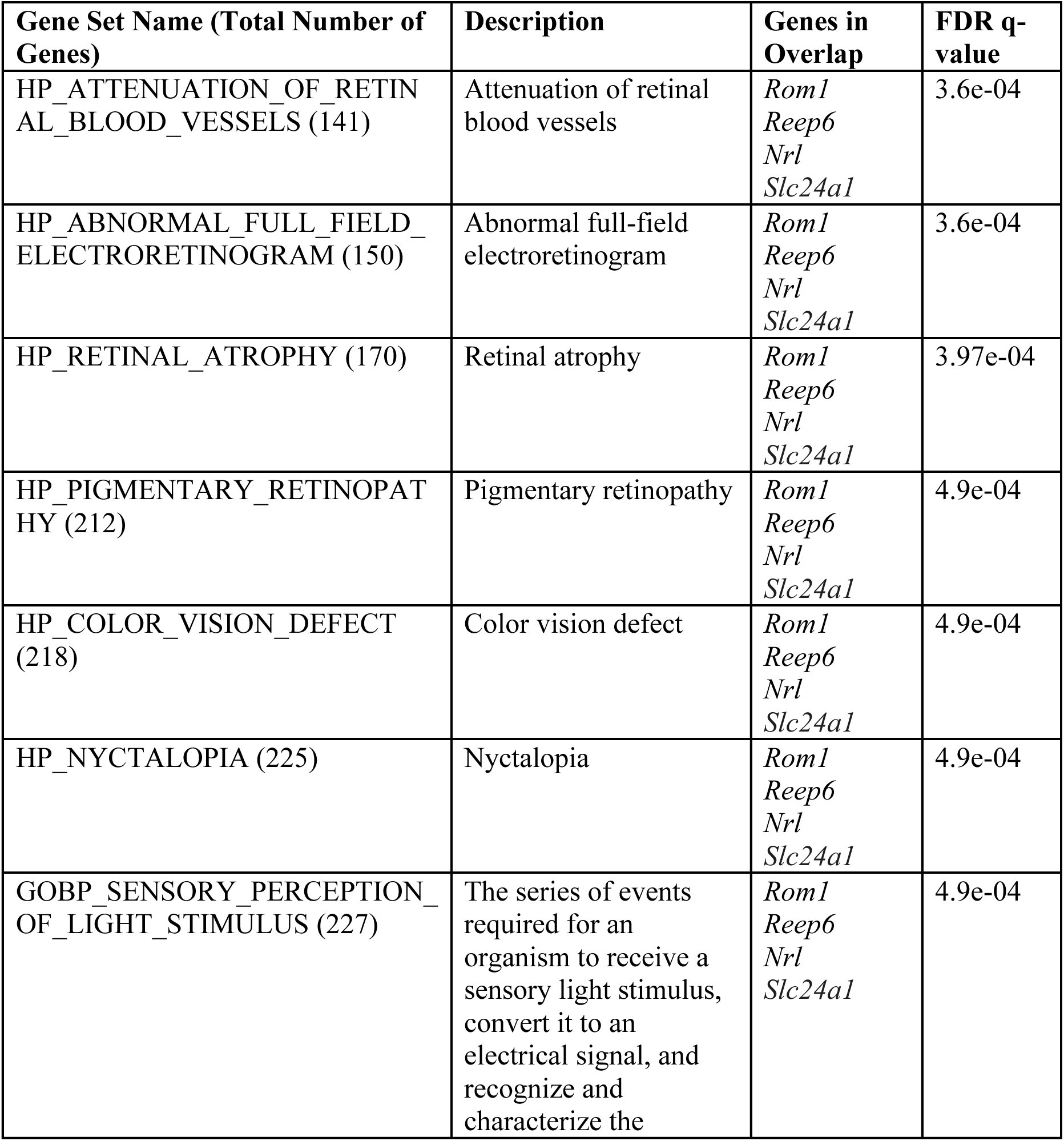
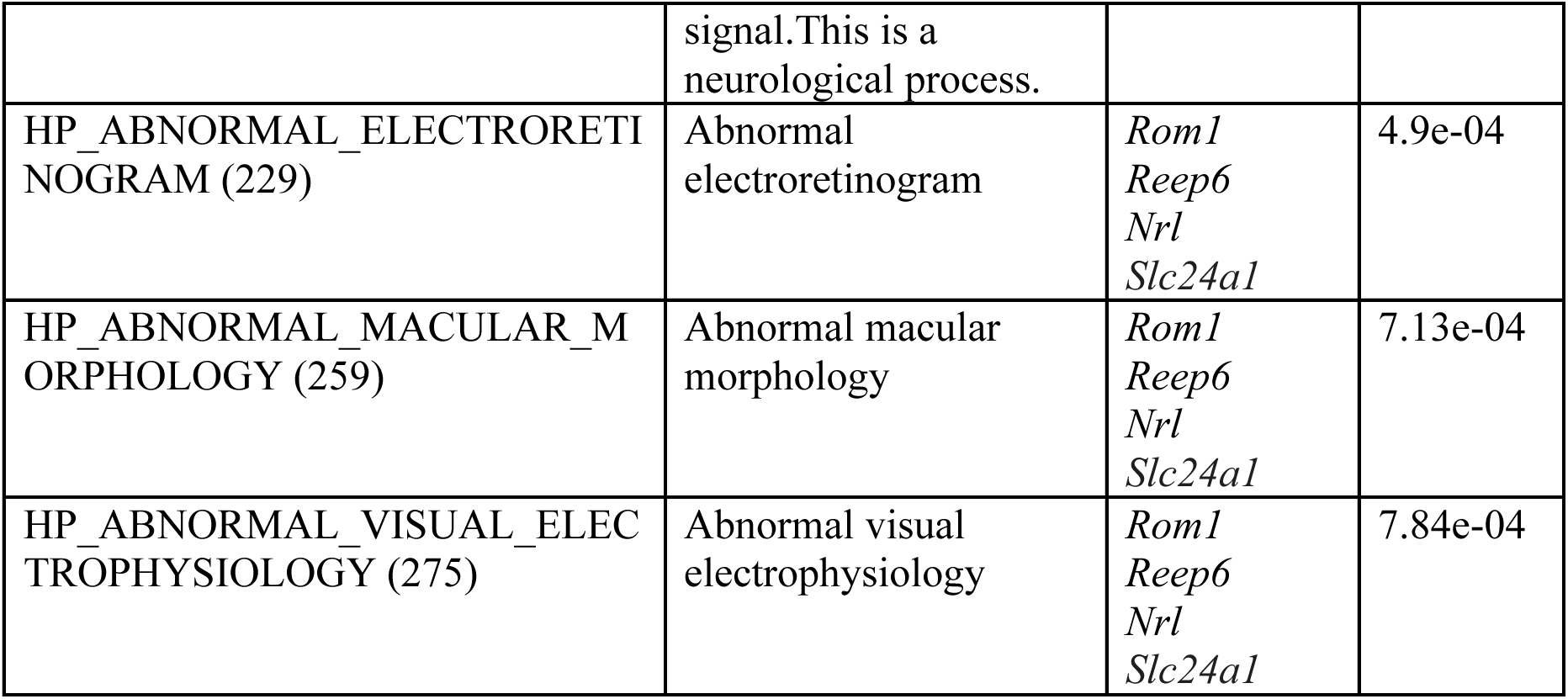
shows the top 10 pathways and gene sets significantly enriched by the genes most predictive of TUNEL apoptosis phenotype, using KEGG and Gene Ontology databases.

| Gene Set Name (Total Number of Genes) | Description | Genes in Overlap | FDR q-value |
| --- | --- | --- | --- |
| HP_ATTENUATION_OF_RETINAL_BLOOD_VESSELS (141) | Attenuation of retinal blood vessels | *Rom1*<br>*Reep6*<br>*Nrl*<br>*Slc24a1* | 3.6e-04 |
| HP_ABNORMAL_FULL_FIELD_ELECTRORETINOGRAM (150) | Abnormal full-field electroretinogram | *Rom1*<br>*Reep6*<br>*Nrl*<br>*Slc24a1* | 3.6e-04 |
| HP_RETINAL_ATROPHY (170) | Retinal atrophy | *Rom1*<br>*Reep6*<br>*Nrl*<br>*Slc24a1* | 3.97e-04 |
| HP_PIGMENTARY_RETINOPATHY (212) | Pigmentary retinopathy | *Rom1*<br>*Reep6*<br>*Nrl*<br>*Slc24a1* | 4.9e-04 |
| HP_COLOR_VISION_DEFECT (218) | Color vision defect | *Rom1*<br>*Reep6*<br>*Nrl*<br>*Slc24a1* | 4.9e-04 |
| HP_NYCTALOPIA (225) | Nyctalopia | *Rom1*<br>*Reep6*<br>*Nrl*<br>*Slc24a1* | 4.9e-04 |
| GOBP_SENSORY_PERCEPTION_OF_LIGHT_STIMULUS (227) | The series of events required for an organism to receive a sensory light stimulus, convert it to an electrical signal, and recognize and characterize the signal. This is a neurological process. | *Rom1*<br>*Reep6*<br>*Nrl*<br>*Slc24a1* | 4.9e-04 |
| HP_ABNORMAL_ELECTRORETINOGRAM (229) | Abnormal electroretinogram | *Rom1*<br>*Reep6*<br>*Nrl*<br>*Slc24a1* | 4.9e-04 |
| HP_ABNORMAL_MACULAR_MORPHOLOGY (259) | Abnormal macular morphology | *Rom1*<br>*Reep6*<br>*Nrl*<br>*Slc24a1* | 7.13e-04 |
| HP_ABNORMAL_VISUAL_ELECTROPHYSIOLOGY (275) | Abnormal visual electrophysiology | *Rom1*<br>*Reep6*<br>*Nrl*<br>*Slc24a1* | 7.84e-04 |

### Comparison to results from CRISP ensemble for TUNEL

Because CRISP is a binary classifier and the TUNEL values are continuous, we binned the TUNEL phenotype values as “low” and “high” based on their being lower or higher than the median TUNEL value, respectively. The top 20 genes most predictive of the TUNEL phenotype according to the CRISP ensemble that were also identified by our approach include *Mgarp*, *Nrl*, *Reep6*, *Slc24a1*, and *Vtn*. Moreover, when submitting the 20 CRISP genes to MSigDB, the gene sets that intersect with those enriched by our approach include GOBP_SENSORY_PERCEPTION_OF_LIGHT_STIMULUS, HP_ABNORMAL_FULL_FIELD_ELECTRORETINOGRAM, HP_ABNORMAL_MACULAR_MORPHOLOGY, HP_ABNORMAL_VISUAL_ELECTROPHYSIOLOGY, and HP_ATTENUATION_OF_RETINAL_BLOOD_VESSELS.

## Discussion

In this section, we highlight our observations of spaceflight effects on ocular physiology and molecular pathways. We discuss the results of our machine learning experiments that predict 4-HNE and TUNEL values from gene expression, and we conclude with strengths, limitations, and future directions.

### 4-HNE Genes and Pathways

The most predictive genes for the 4-HNE phenotype (Table 2) include *B2m*, *Uqcr10*, *Tf*, *Arl6ip1*, *Cnga1*, *mt-Nd1*, *Pcare*, *Hnrnph1*, *Snap25*, *Hnrnpa2b1*, *Cplx1*, *Atf4*, *Syn1*, *Hsp90ab1*, *Sncb*, *Nr2e3*, *Cox7b*, and *Efemp1*. Enriched Gene Ontology and Human Phenotype Ontology pathways include optic disc pallor, ophthalmoparesis, retinal pigmentation abnormalities, abnormal optic nerve morphology, retinal dystrophy, glaucoma, and monoatomic cation pathways.

Genes *B2m*, *Tf*, *Arl6ip1*, and *Uqcr10* converge on membrane-associated pathways (GO:0098796). Upregulated *B2m* reflects heightened immune signaling following oxidative injury^35^. *Tf* expression changes suggest altered iron homeostasis under oxidative stress^36^. *Arl6ip1*, an apoptotic regulator, supports adaptive responses to endoplasmic reticulum and membrane stress through mitochondria-associated membrane localization^37^. Elevated *Uqcr10* reflects compensatory mitochondrial remodeling since 4-HNE disrupts respiratory-chain proteins^38^. 4-HNE also modifies essential photoreceptor proteins. *Cnga1* disruption impairs cation flow and signal conversion^39^, *mt-Nd1* modification reduces ATP and elevates ROS^40^, and *Pcare* disruption impairs photoreceptor disc organization^41^. *Hnrnph1* enrichment indicates 4-HNE affects RNA processing supporting retinal and optic nerve function^42^, while *Snap25* and *Cplx1* enrichment reflect synaptic protein susceptibility to lipid peroxidation and contributions to optic nerve deterioration^43–45^. *Efemp1* dysregulation causes Doyne honeycomb retinal dystrophy, myopia, glaucoma, and drusen accumulation^46^. Spaceflight-disrupted integrin signaling affects retinal vascular permeability, while microgravity-induced collagen stretching^47^ potentially causes SANS eye deformation. 4-HNE directly impacts ERK levels^48^, increasing cell stress. Finally, *Atf4*, *Syn1*, *Hsp90Ab1*, and *Sncb* relate to neuron projection and cation transport: *Atf4* regulates vascular apoptosis^49^, *Hsp90Ab1* prevents ER stress-induced apoptosis^50^, 4-HNE alters *Syn1* phosphorylation disrupting vesicle tethering^51^, and *Sncb* modification promotes oxidative stress and inflammation^52^, with chronic 4-HNE elevation ultimately causing synaptic dysfunction.

### TUNEL Genes and Pathways

The TUNEL phenotype analysis highlighted key predictive genes: *Ddit4*, *Nrl*, *Tcf20*, *Pdzph1*, *Gabarapl1*, *Ip6k2*, *Mgarp*, *Reep6*, *Rom1*, *Slc24a1*, and *Vtn*, with enrichment in apoptotic signaling, photoreceptor cell death, structural deterioration, and retinopathy pathways. These results are consistent with TUNEL’s detection of DNA fragmentation indicating apoptosis. Endothelial dysfunction is particularly relevant, as Zanello et al. found corneal bullae in spaceflown mice^53^, and retinal vascular endothelial dysfunction may link to cotton wool spots seen in astronauts, consistent with blood-retinal barrier disruption documented in RR-9 mission samples^8^.

*Ddit4* regulates stress-induced apoptosis through mTORC1 inhibition and ROS control^54^, with upregulation activating autophagy and apoptosis via the mTOR axis under neurotoxic and hypoxic conditions^55–57^. This directly parallels microgravity-induced stress responses in spaceflight conditions. *Nrl* is critical for rod photoreceptor differentiation and homeostasis, with inhibition preventing cell death across multiple retinal degeneration models^58^. It regulates rod-specific genes including *Rom1*, *Nr2e3*, and *Cngb1*^59^. *Rom1* disruption causes progressive photoreceptor degeneration and outer nuclear layer thinning^60^, consistent with spaceflight-induced photoreceptor dysfunction observed experimentally^10^. *Reep6* maintains rod photoreceptor ER homeostasis and phototransduction protein trafficking, with deficiency causing early-onset dysfunction through ER stress^61^. *Rom1* is required for rod photoreceptor viability and the regulation of disk morphogenesis^62^, while *Pde6g* dysregulation further compromises the phototransduction cascade^63^. Dysregulation of either contributes to photoreceptor degeneration. *Gabarapl1*, essential for autophagosome-lysosome fusion^64^, shows elevated expression under retinal oxidative stress with dual protective roles acutely, though chronic elevation may transition to pathological autophagy^65,66^. *Tcf20* functions as a transcriptional coactivator with widespread tissue expression^67,68^, and *Pdzph1* likely reflects compromised protein trafficking or synaptic integrity given PDZ domain proteins’ known roles in photoreceptor-bipolar cell connections^69^. *Vtn* promotes retinal neurite outgrowth and cell adhesion through integrin-RGD interactions^70,71^, and reduced expression could impair photoreceptor-epithelial interactions under spaceflight stress. Finally, *Mgarp*, highly expressed in photoreceptor inner segments^72^, regulates mitochondrial transport under hypoxic stress^73^, and its reduced expression may compromise mitochondrial trafficking to photoreceptor terminals during spaceflight.

## Conclusion

This machine learning ensemble identified distinct gene signatures predictive of 4-HNE-mediated oxidative damage and TUNEL-positive apoptosis in spaceflown murine retinae. These genes implicate membrane dysfunction, photoreceptor degeneration, and apoptotic signaling as complementary pathological mechanisms in SANS. Together, these findings provide a molecular framework for identifying noninvasive biomarkers and therapeutic targets to monitor and protect astronaut visual health during long-duration deep-space missions. As humanity extends its presence beyond low Earth orbit, such molecular insights and translational strategies will prove essential for safeguarding crew health and mission success during deep-space exploration.

## Strengths and Limitations

4-HNE immunostaining directly indicates lipid peroxidation with stable adducts that facilitate post-flight analysis despite sample transport delays, enabling localized tissue visualization. Limitations include lack of SANS specificity, potential cross-reactivity, inter-laboratory ELISA variability, and uncertainty regarding adduct detection timing post-exposure^74^. TUNEL provides sensitive, spatially resolved histological readout of DNA fragmentation across vulnerable retinal layers including GCL, INL, and photoreceptors^75^. However, it cannot reliably distinguish apoptosis from necrosis, autolysis, or irradiation-induced damage^76^, reflects only late-stage cell death, and sensitivity varies with tissue handling and post-flight processing delays^77^. Computationally, the small sample size (n=16) limits generalizability, and identified genes represent associations rather than confirmed causal mechanisms. However, because our approach yielded results consistent with those generated by the CRISP causal inference classification platform, we may infer that our associations are robust to spurious correlations. Bulk RNA-sequencing provides quantifiable molecular visibility to cell response to spaceflight, but it’s only a single point-in-time snapshot which averages expression across heterogeneous cell populations. While our genes were found to be very informative to our model predictions, gene filtering likely excluded regulatory RNAs or low-abundance transcription factors with potentially important mechanistic roles.

## Future Directions

The genes and pathways we identified as predictive of the 4-HNE and TUNEL phenotypes require experimental validation *in vitro* and *in vivo*. Temporal profiling across preflight, during-flight, and post-flight timepoints would capture molecular response dynamics, following the NASA Twins Study and Inspiration 4 paradigms^78,79^. Single-cell and spatial transcriptomics would resolve cell-type-specific vulnerabilities across photoreceptors, endothelial cells, and ganglion cells^80^. Expanded sample sizes across multiple missions would strengthen confidence in our findings. Human ground-based analogs including head-down tilt bed rest^81^, dry immersion^82^, and prospective astronaut studies using accessible biospecimens or imaging biomarkers would establish clinical relevance. Pupillometry represents a promising noninvasive biomarker for intracranial pressure changes during spaceflight^83^, and novel visual assessment technologies continue advancing understanding of SANS pathophysiology^84^. More broadly, this work exemplifies machine learning’s growing role in extracting actionable insights from spaceflight datasets to support autonomous biomonitoring^85^, self-driving laboratory capabilities^86^, and crew health monitoring for lunar and Martian exploration.

## Acknowledgements

The authors would like to thank the NASA Biological and Physical Sciences Division for supporting NASA Ames Life Sciences Data Archive, NASA GeneLab, and its umbrella project the NASA Open Science Data Repository. While this paper was not completely organized as one of the open projects of the OSDR-Analysis Working Groups, a number of the authors are AWG members and credit is due. We are grateful for all discussions and contributions especially from the Causal Inference Subgroup of the OSDR-AI/ML AWG. RTS and JAC are both part of the “AI for Life in Space” group of researchers at NASA Ames.

## Data and Code Availability

All experimental data used in this study are publicly available through NASA’s Open Science Data Repository, the expansion of NASA GeneLab to integrate the NASA Ames Life Sciences Data Archive to include high quality physiological, phenotypic, bioimaging, environmental telemetry, and behavior (https://science.nasa.gov/biological-physical/data/osdr/). The three Rodent Research 9 (RR-9) mission datasets were mined from: 1) OSD-255 containing retinal RNA sequencing gene expression data (https://doi.org/10.26030/mebr-1747), 2) OSD-557 containing 4-HNE immunostaining microscopy data for oxidative stress quantification (https://doi.org/10.26030/yv31-1a54), and 3) OSD-568 containing TUNEL assay microscopy data for apoptosis detection (https://doi.org/10.26030/d09k-4e68). Gene set enrichment analyses were performed using the Molecular Signatures Database (MSigDB; https://www.gsea-msigdb.org/gsea/msigdb). All Python code for data preprocessing, machine learning model training, feature selection, and visualization is available at https://colab.research.google.com/drive/11moJztlWNqPYNUfef6Ids9mOl1xmpSXb?usp=sharing.

## Authors’ Contributions

LMS, JAC, RTS, and PAV were co-investigators in the grant described in the Funding section. This grant led to the design and development of an online training course in which high school students learned how to use artificial intelligence and machine learning to study the effect of spaceflight on living systems. JAC designed and implemented the Jupyter notebooks for that course to analyze the molecular retina data from the RR9 mission. The students in that course wrote reports based on the results of these analyses. JAC developed the first version of this manuscript by integrating components from the reports of the students who expressed interest in participating in this research (AR, Aarav J, AC, AA, AP, AN, Anishka J, AB, AK, Anagha J, BP, DS, DN, EK, EP, FPU, IS, IY, JW, JL, KS, MPDM, MH, ML, MV, Misha K, Mrinalini K, MAJ, MA, NZC, NA, NBC, RH, RP, SD, Samarth S, Shawnak S, SG, Shriya S, SC, VP, VN, and ZS). PP and JC organized the efforts of the students. DMT and AL ran the CRISP experiments. The biomedical experts (JO, XWM, SAN, EW, and LMS) provided feedback and guidance for revising the first version of the manuscript. RTS and JAC wrote the final version of the manuscript based on that feedback. PAV, SG, and JMG provided management oversight and support of the research. All authors proofread the manuscript and provided their feedback.

