## Supplement for "Machine Learning Ensemble Reveals Distinct Molecular Pathways of Retinal Damage in Spaceflown Mice"

### Appendix

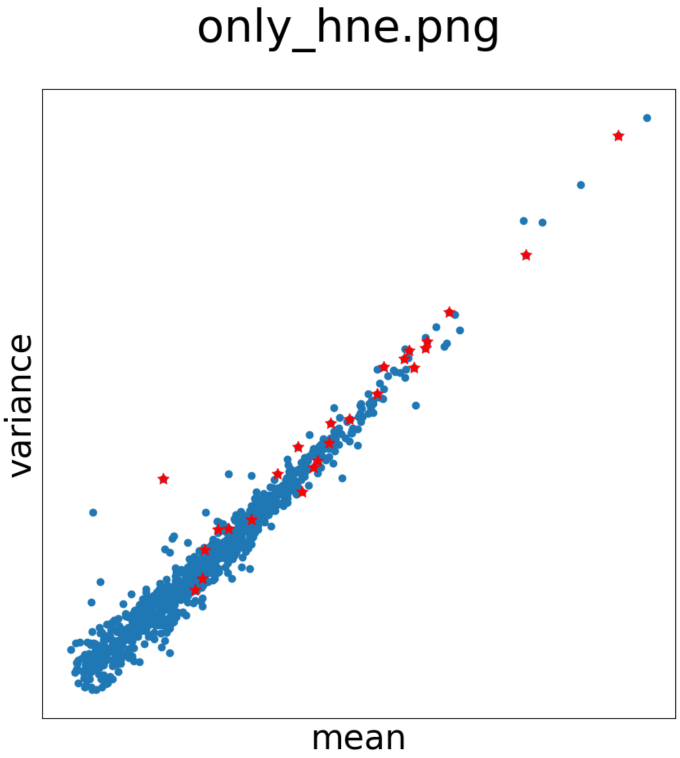


**Figure A1**. Distribution of genes identified as most predictive only of the HNE phenotype by model (in red) against background of all genes used as features to build HNE model (in blue).

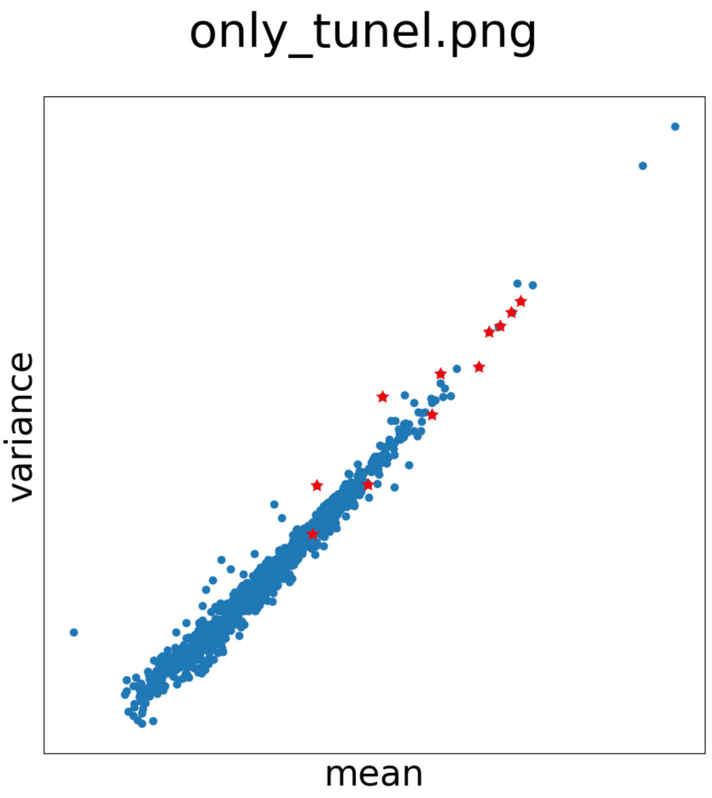


**Figure A2**. Distribution of genes identified as most predictive of TUNEL by model ensemble (in red) against background of all genes used as features to build TUNEL model (in blue).

#### Genes most predictive of both HNE and TUNEL phenotypes

We took the intersection of the HNE and TUNEL results to determine which genes are implicated in both of these and potentially other phenotypes which are characteristic of SANS. Table 6 lists the intersection of genes from the HNE and TUNEL results.

| |
|---|
| *Atp5f1a Dynlrb1 Prph2 Rlbp1 Stmn3* |

**Table A1**. Intersection of genes from the TUNEL and HNE majority result sets.

The top genes most predictive of both the HNE oxidative stress phenotype and the TUNEL apoptosis phenotype significantly enrich the gene sets listed in Table 7.

| **Gene set name (number of genes)** | **description** | **# genes in overlap** | **FDR valu** |
|---|---|---|---|
| HP_PROGRESSIVE_VISUAL_FIELD_DEFECTS (7) | Progressive visual field defects | *Prph2 Rlbp1* | 2.37e |

| | | | |
|---|---|---|---|
| HP_RETINAL_FLECKS (9) | Retinal flecks | *Prph2*<br>*Rlbp1* | 2.37e |
| HP_LENTICONUS (10) | Lenticonus | *Prph2*<br>*Rlbp1* | 2.37e |
| HP_ABNORMAL_FOVEAL_MORPHOLOGY_ON_MACULAR_OCT (11) | Abnormal foveal morphology on macular OCT | *Prph2*<br>*Rlbp1* | 2.37e |
| HP_ROD_CONE_DYSTROPHY (236) | Rod-cone dystrophy | *Prph2*<br>*Rlbp1*<br>*Atp5f1a* | 4.97e |
| HP_ABSENT_FOVEAL_REFLEX (20) | Absent foveal reflex | *Prph2*<br>*Rlbp1* | 4.97e |
| HP_CONGENITAL_STATIONARY_NIGHT_BLINDNESS (21) | Congenital stationary night blindness | *Prph2*<br>*Rlbp1* | 4.97e |
| HP_ABNORMALITY_OF_LENS_SHAPE (22) | Abnormality of lens shape | *Prph2*<br>*Rlbp1* | 4.97e |
| HP_BLINDNESS (297) | Blindness | *Prph2*<br>*Rlbp1*<br>*Atp5f1a* | 5.51e |
| HP_OPHTHALMOPARESIS (305) | Ophthalmoparesis | *Prph2*<br>*Rlbp1*<br>*Atp5f1a* | 5.51e |

**Table A1** shows the top 10 pathways significantly enriched for the genes most predictive of both HNE oxidative stress and TUNEL apoptosis phenotypes, using KEGG and Gene Ontology databases.

Taken together, the HNE and TUNEL stain results suggest that oxidative stress and apoptotic cell death may be a combinatorial mechanism causing retinal damage due to microgravity. The overlap concerning transport (*Atp5f1a, Dynlrb1, Stmn3*) and protein binding (*Prph2, Rlbp1*) confirms the critical role of photoreceptor bioenergetics and structure in susceptibility to spaceflight-induced damage. A molecular mechanism may exist whereby mitochondrial oxidative damage to photoreceptors results in apoptotic cell death. This mechanism is similar to what is currently thought to occur within SANS and confirms that the molecular mechanisms elucidated within this study are relevant to this human symptomology.
